# Analysis of the immunological biomarker profile during acute Zika virus infection reveals the overexpression of CXCL10, a chemokine already linked to neuronal damage

**DOI:** 10.1101/185041

**Authors:** Felipe Gomes Naveca, Gemilson Soares Pontes, Aileen Yu-hen Chang, George Allan Villarouco da Silva, Valdinete Alves do Nascimento, Dana Cristina da Silva Monteiro, Marineide Souza da Silva, Lígia Fernandes Abdalla, João Hugo Abdalla Santos, Tatiana Amaral Pires de Almeida, Matilde del Carmen Contreras Mejía, Tirza Gabrielle Ramos de Mesquita, Helia Valeria de Souza Encarnação, Matheus de Souza Gomes, Laurence Rodrigues Amaral, Ana Carolina Campi-Azevedo, Jordana Graziela Coelho-dos-Reis, Lis Ribeiro do Vale Antonelli, Andréa Teixeira-Carvalho, Olindo Assis Martins-Filho, Rajendranath Ramasawmy

**Affiliations:** Programa de Pós-Graduação em Biologia da Interação Patógeno-Hospedeiro, Instituto Leônidas e Maria Deane - Fiocruz Amazônia, Manaus, Amazonas, Brazil; Programa de Pós-Graduação em Imunologia Básica e Aplicada, Universidade Federal do Amazonas, Manaus, Amazonas, Brazil; Instituto Nacional de Pesquisas da Amazônia, Manaus, Amazonas, Brazil; George Washington University, Washington, DC, United States of America; Universidade do Estado do Amazonas, Manaus, Amazonas, Brazil; Hospital Adventista de Manaus, Manaus, Amazonas, Brazil; Fundação de Medicina Tropical - Dr Heitor Vieira Dourado, Manaus, Amazonas, Brazil; Universidade Federal de Uberlândia, Patos de Minas, Minas Gerais, Brazil; Centro de Pesquisas René Rachou - Fiocruz Minas, Belo Horizonte, Minas Gerais, Brazil; Universidade Nilton Lins, Manaus, Amazonas, Brazil

**Author notes:** (FGN). (OAMF). These authors contributed equally to this work (Felipe Gomes Naveca, Olindo Assis Martins-Filho). These authors also contributed equally to this work (Gemilson Soares Pontes, Aileen Yu-hen Chang, Andréa Teixeira-Carvalho).

## Abstract

Infection with Zika virus (ZIKV) manifests in a broad spectrum of disease ranging from mild illness to severe neurological complications. To define immunologic correlates of ZIKV infection, we characterized the levels of circulating cytokines, chemokines and growth factors in 54 infected patients of both genders, at five different time-points after symptoms onset using microbeads multiplex immunoassay; statistical analysis and data mining compared to 100 age-matched controls. ZIKV-infected patients present a striking systemic inflammatory response with high levels of pro-inflammatory mediators. Despite the strong inflammatory pattern, IL-1Ra and IL-4 are also induced during acute infection. Interestingly, the inflammatory cytokines, IL-1β, IL-13, IL-17, TNF-α, IFN-γ; chemokines, CXCL8, CCL2, CCL5; and the growth factor G-CSF display a bimodal distribution accompanying viremia. While this is the first manuscript to document bimodal distributions of viremia in ZIKV infection, bimodal viremia has been documented in other viral infections with primary viremia peaks during mild systemic disease and a secondary viremia with distribution of the virus to organs and tissues. Moreover, biomarker network analysis demonstrated distinct dynamics in consonance with the bimodal viremia profiles at different time-points during ZIKV infection. Such robust cytokine and chemokine response has been associated with blood-brain barrier permeability and neuroinvasiveness in other flaviviral infections. High-dimensional data analysis further established CXCL10, a chemokine involved in fetal neuron apoptosis and Guillain-Barré syndrome, as the most promising biomarker of acute ZIKV infection for a potential clinical application.

**Author Summary:** Infection with Zika virus manifests in a broad spectrum of disease ranging from mild illness to severe neurological complications. This study characterized the levels of circulating cytokines, chemokines and growth factors in Zika-infected patients showing an inflammatory immune response. Specifically, this study identified a chemokine, CXCL10, known to be involved in fetal neuron apoptosis and Guillain-Barré syndrome, as the most promising biomarker to characterize acute Zika virus infection.

## Introduction

The Zika virus (ZIKV) is an arthropod-borne *Flavivirus*, transmitted mainly by the bite of female *Aedes* mosquitos, that usually causes a mild illness characterized by conjunctivitis, pruritus, muscle and joint pain, rash and slight fever [1]. Outbreaks of ZIKV infection were first recorded in Micronesia and later in French Polynesia, where atypical manifestations were initially documented, including the Guillain-Barré Syndrome [2,3]. In Brazil, ZIKV infection during pregnancy was linked to an unusual increase in the number of microcephaly cases [4]. Following the Brazilian report of congenital malformations, the number of microcephaly cases in French Polynesia were reanalyzed, and a connection with ZIKV was further established [5]. The broad spectrum of fetal clinical manifestations resulting from ZIKV infection lead to a new classification termed Zika Congenital Syndrome [6].

Host immune response plays an important role in the clinical course of patients with viral infection. Particularly, cellular immunity and key components of the innate immune response, such as interferons and other cytokines/chemokines, play an essential role in limiting the viral spread [7]. To date, only two studies describing immune mediators in Zika-infected patients have been reported [8,9]. In Tappe et al, reliable immunological biomarker profile during acute infection could not be established due to the small sample size. Kam et al. describes immune markers from a cohort from Campinas, Brazil showing inflammatory immune response and several immune mediators specifically higher in ZIKV-infected patients, with levels of CXCL10, IL-10, and HGF differentiating between patients with and without neurological complications. Kam et al. also found higher levels of CXCL10, IL-22, MCP-1, and TNF-α were observed in ZIKV-infected pregnant women carrying babies with fetal growth associated malformations.

In this study, we evaluated the immune response during the acute ZIKV infection by analysis the serum levels of cytokines, chemokines and growth factors from an adult cohort from Manaus, Brazil of 54 ZIKV-infected cases and 100 controls over five time points during symptomatic ZIKV infection. We present the time course of cytokine response in relation to viremia and identify a chemokine that may serve as a biomarker of acute ZIKV infection, providing new insights into ZIKV neuropathogenesis.

## Methods

### Study Population and Design

We used a non-probabilistic convenience sampling and a cross-sectional experimental design, together with robust statistical analysis and data mining, for the evaluation of the immunological biomarker profile during acute ZIKV infection. In the first semester of 2016, a total of 54 suspected ZIKV-infected cases (29 non-pregnant females and 25 males, all adults) were recruited at Hospital Adventista de Manaus, Amazonas state, Brazil. All patients presented a maculopapular rash, with or without fever, and at least one of the following symptoms: pruritus, arthralgia, joint swelling or conjunctival hyperemia, within five days after the symptoms onset. Age-matched non-infected (NI) controls, females (46) and males (54), were enrolled for comparison and basic characteristics, including physical examination and virological findings are provided. Comprehensive laboratory records were available for 21 patients (15 male and six female), including routine laboratory tests.

### Ethics Statement

The study protocol was approved by the Ethics Committee of the Universidade do Estado do Amazonas (CAAE: 56745116.6.0000.5016) and all subjects included provided written informed consent.

### Differential molecular diagnosis of Zika and viral load estimative

Serum samples were sent to Fiocruz Amazônia and tested for ZIKV (envelope coding region) [10], Chikungunya (CHIKV) [11] and Dengue (DENV) [12] by RT-qPCR. Samples positive for CHIKV or DENV were excluded. Sample inclusion criteria also required the internal control (spiked MS2 bacteriophage) to display a Ct value between 30-32. The viremia was indirectly estimated by RT-qPCR and reported as 1/Ct*100.

### Dengue virus serology

Serum samples were tested for previous exposure to DENV with Serion ELISA classic Dengue Virus IgG (Institut Virion/Serion GmbH, Germany).

### Microbeads assay for serum biomarkers

High-performance microbeads 27-plex assay (Bio-Rad, Hercules, CA, USA) was employed for detection and quantification of multiple targets, including: CXCL8 (IL-8); CXCL10 (IP-10); CCL11 (Eotaxin); CCL3 (MIP-1α); CCL4 (MIP-1β); CCL2 (MCP-1); CCL5 (RANTES); IL-1β, IL-6, TNF-α; IL-12; IFN-γ, IL-17; IL-1Ra (IL-1 receptor antagonist); IL-2; IL-4; IL-5; IL-7; IL-9; IL-10; IL-13; IL-15; FGF-basic; PDGF; VEGF; G-CSF and GM-CSF. Samples were tested according to the manufacturer’s instructions on a Bio-Plax 200 instrument (Bio-Rad). The serum levels of IL-2, IL-7, and IL-15 were below the detection limits in several samples and were excluded of further analysis. The results were expressed as pg/mL.

### Statistical analysis and Data mining

Statistical analyses were initially performed using GraphPad Prism (GraphPad Software 6.0, San Diego, CA, USA). Outliers within each measurement group were identified by the ROUT Method (Q=1%) and removed. Cleaned data was then used for the evaluation of Gaussian distribution with D’Agostino & Pearson omnibus normality test.

Comparative analysis of the clinical records was carried out by Fisher’s exact test. The analysis of biomarker levels between NI controls *vs.* ZIKV-infected cases, and between genders, was performed by Mann-Whitney test. Multivariate correlations for biomarker levels and routine laboratory tests were analyzed with the nonparametric Spearman’s test (alpha 0.05) running on the JMP Software, v13.1.0 (SAS Institute, Cary, NC, USA). Correlations (Spearman ρ) were represented by a color map matrix.

The dynamics of viremia, chemokines, cytokines and growth factors were evaluated using the median value of each analyte. Comparative analysis of the biomarkers was carried out by Kruskal-Wallis followed by Dunn’s post-test. For all tests, significant differences were considered at two-tailed p<0.05.

Data management strategies were applied to identify general and time-specific profiles. Biomarker signature analysis was carried out as previously described [13]. Radar charts were assembled to compile the biomarker signature of NI controls and ZIKV-infected cases applying the 75^th^ percentile as threshold. Venn diagram scrutiny was carried out to identify attributes, along with the timeline of the symptoms onset http://bioinformatics.psb.ugent.be/webtools/Venn/. Cytoscape software v3.2.0 (http://www.cytoscape.org/) was employed for visualizing and integrating multiple attributes into circular nodal networks. Connecting edges were drawn to underscore the association as positive (solid line) or negative (dashed line). The biomarker cluster pattern was defined by heatmaps assembled using R software (heatmap.2 function; v3.0.1). Decision tree algorithms were generated with WEKA software v3.6.11 (University of Waikato, New Zealand) to identify root and branch attributes, segregating patients from controls. ROC curves were built to define the cut-off and biomarkers with better performance to discriminate ZIKV-infected patients from NI controls. Performance indices (co-positivity, co-negativity, positive and negative likelihood ratio) were calculated using the MedCalc software v7.3 (Ostend, Belgium).

## Results

### Demographics, clinical records and virological data

The 54 Brazilian Zika cases, 29 non-pregnant females (median age 38 years, IQR 27.5 - 46.5) and 25 males (median age 37 years, IQR 30 - 50), were enrolled between the first and the fifth day after the symptoms onset. A group of 100 non-infected control subjects who were residents of Manaus, Amazonas, Brazil were also included (46 females (median age 28 years, IQR 23 - 36) and 54 males (median age 29.5 years, IQR 23 - 36)). The median viremia expressed as 1/Ct*100 was 2.9 (min=2.7; max=4.2; IQR: 2.8 - 3.0). The frequency of specific ZIKV symptoms was similar between men and woman with the only exception that men had increase frequency of fever compared to woman (100% versus 67%, p=0.005) (Table 1). The DENV IgG testing showed that 94.4% (51/54) of the patients were positive; two had an undetermined result and one male subject was negative.

**Table 1.**
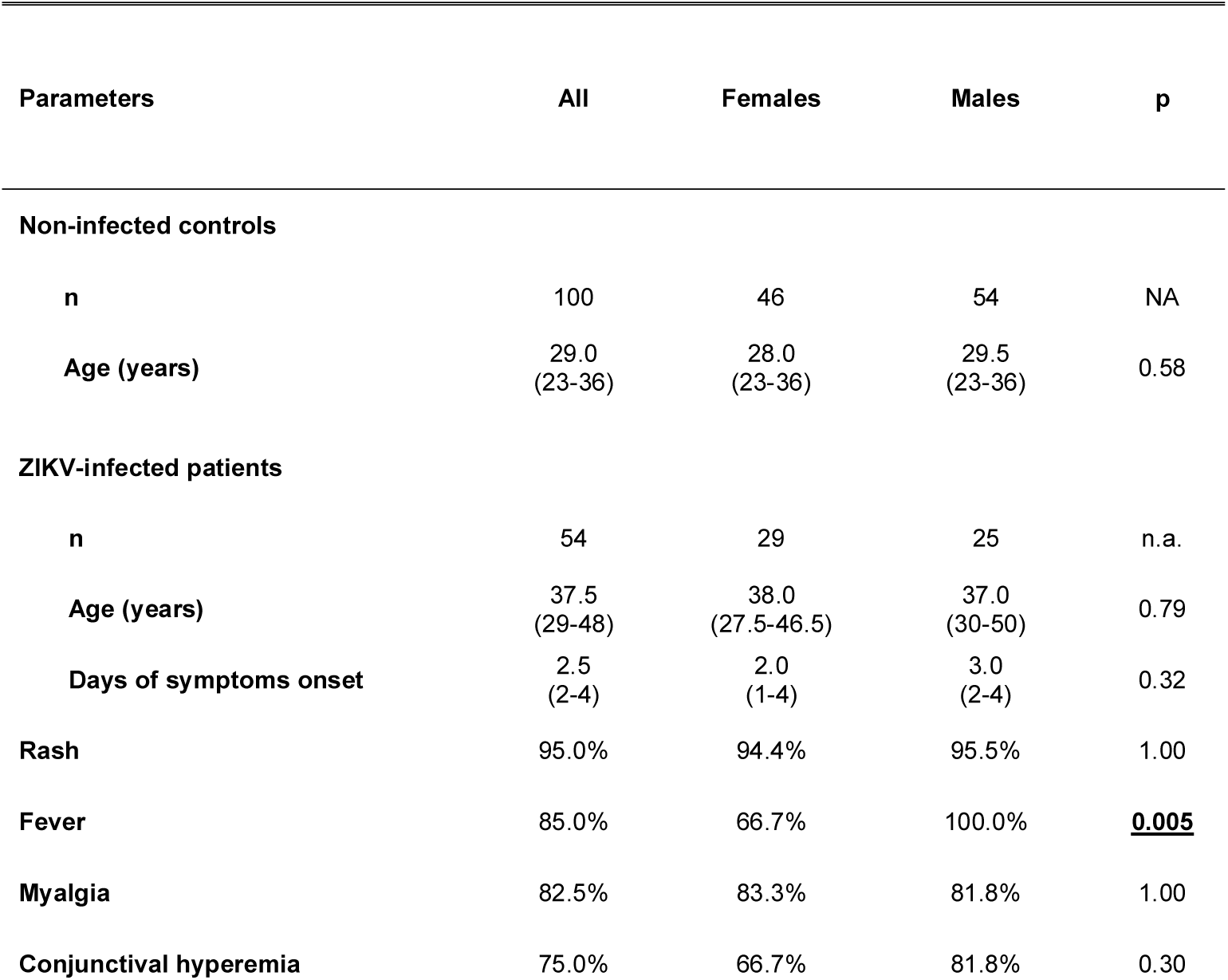
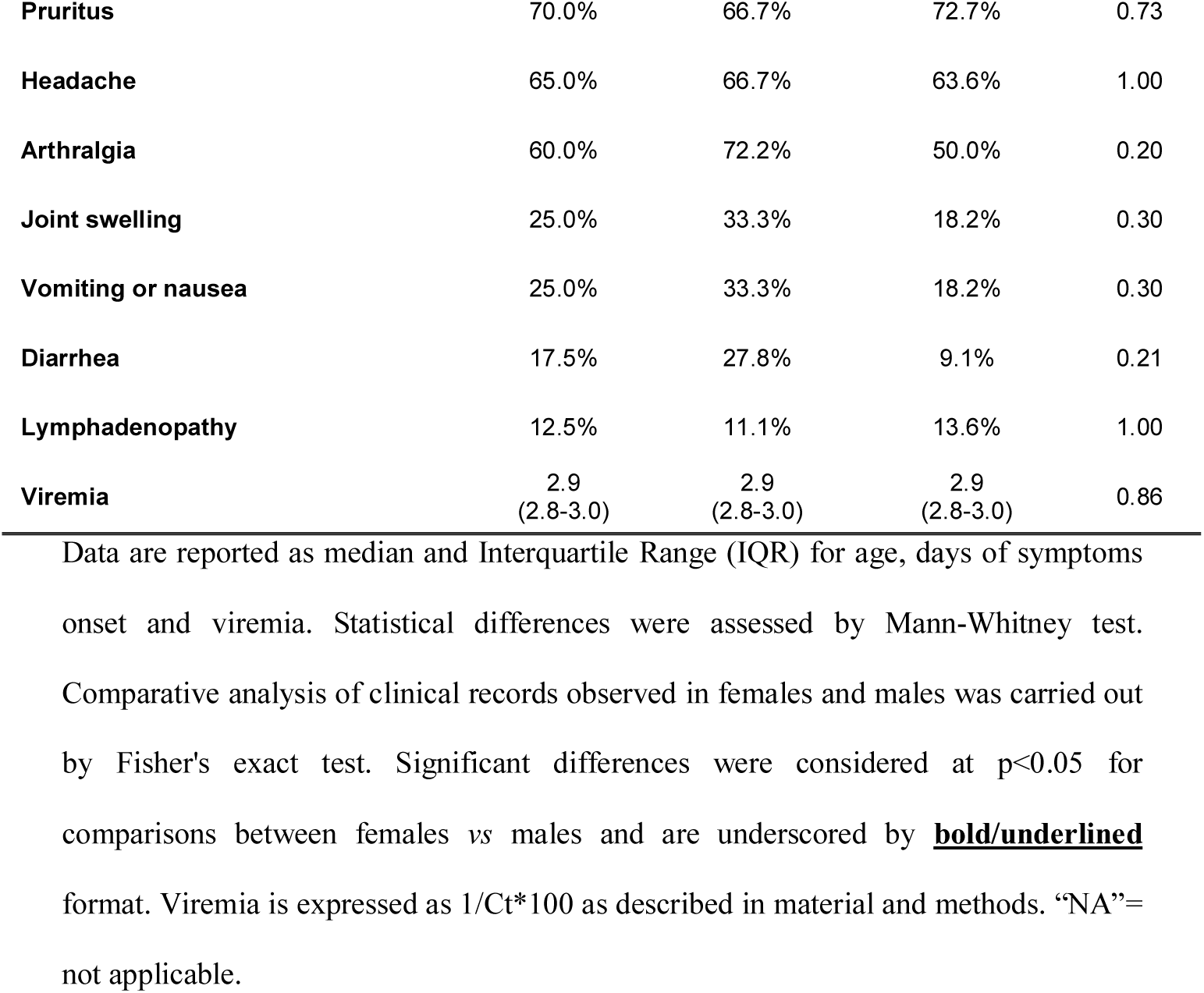
Demographical aspects, clinical records and virological status of ZIKV-infected patients

| Parameters | All | Females | Males | p |
| --- | --- | --- | --- | --- |
| **Non-infected controls** |  |  |  |  |
| n | 100 | 46 | 54 | NA |
| Age (years) | 29.0<br>(23-36) | 28.0<br>(23-36) | 29.5<br>(23-36) | 0.58 |
| **ZIKV-infected patients** |  |  |  |  |
| n | 54 | 29 | 25 | n.a. |
| Age (years) | 37.5<br>(29-48) | 38.0<br>(27.5-46.5) | 37.0<br>(30-50) | 0.79 |
| Days of symptoms onset | 2.5<br>(2-4) | 2.0<br>(1-4) | 3.0<br>(2-4) | 0.32 |
| Rash | 95.0% | 94.4% | 95.5% | 1.00 |
| Fever | 85.0% | 66.7% | 100.0% | **0.005** |
| Myalgia | 82.5% | 83.3% | 81.8% | 1.00 |
| Conjunctival hyperemia | 75.0% | 66.7% | 81.8% | 0.30 |
| **Pruritus** | 70.0% | 66.7% | 72.7% | 0.73 |
| **Headache** | 65.0% | 66.7% | 63.6% | 1.00 |
| **Arthralgia** | 60.0% | 72.2% | 50.0% | 0.20 |
| **Joint swelling** | 25.0% | 33.3% | 18.2% | 0.30 |
| **Vomiting or nausea** | 25.0% | 33.3% | 18.2% | 0.30 |
| **Diarrhea** | 17.5% | 27.8% | 9.1% | 0.21 |
| **Lymphadenopathy** | 12.5% | 11.1% | 13.6% | 1.00 |
| **Viremia** | 2.9<br>(2.8-3.0) | 2.9<br>(2.8-3.0) | 2.9<br>(2.8-3.0) | 0.86 |

Data are reported as median and Interquartile Range (IQR) for age, days of symptoms onset and viremia. Statistical differences were assessed by Mann-Whitney test. Comparative analysis of clinical records observed in females and males was carried out by Fisher's exact test. Significant differences were considered at $p < 0.05$ for comparisons between females vs males and are underscored by **bold/underlined** format. Viremia is expressed as $1/Ct \times 100$ as described in material and methods. "NA"= not applicable.

### Correlation of immunological biomarkers during acute ZIKV infection with routine laboratory tests

The results of 45 continuous variables including immunological biomarkers; routine laboratory tests; age; viremia and symptoms onset were analyzed (Fig 1). Overall, moderate correlations were observed for several variables, whereas the strongest correlations were observed between TNF-α and CCL5 (Spearman ρ 0.8245) and Lymphocytes (%) and Neutrophils (%) (Spearman ρ -0.8084). All results were represented in a color map matrix, where statistically supported associations (p<0.05), regarding the routine laboratorial tests and immunological biomarkers, were highlighted (inserted table in Fig 1).

**Fig 1.**
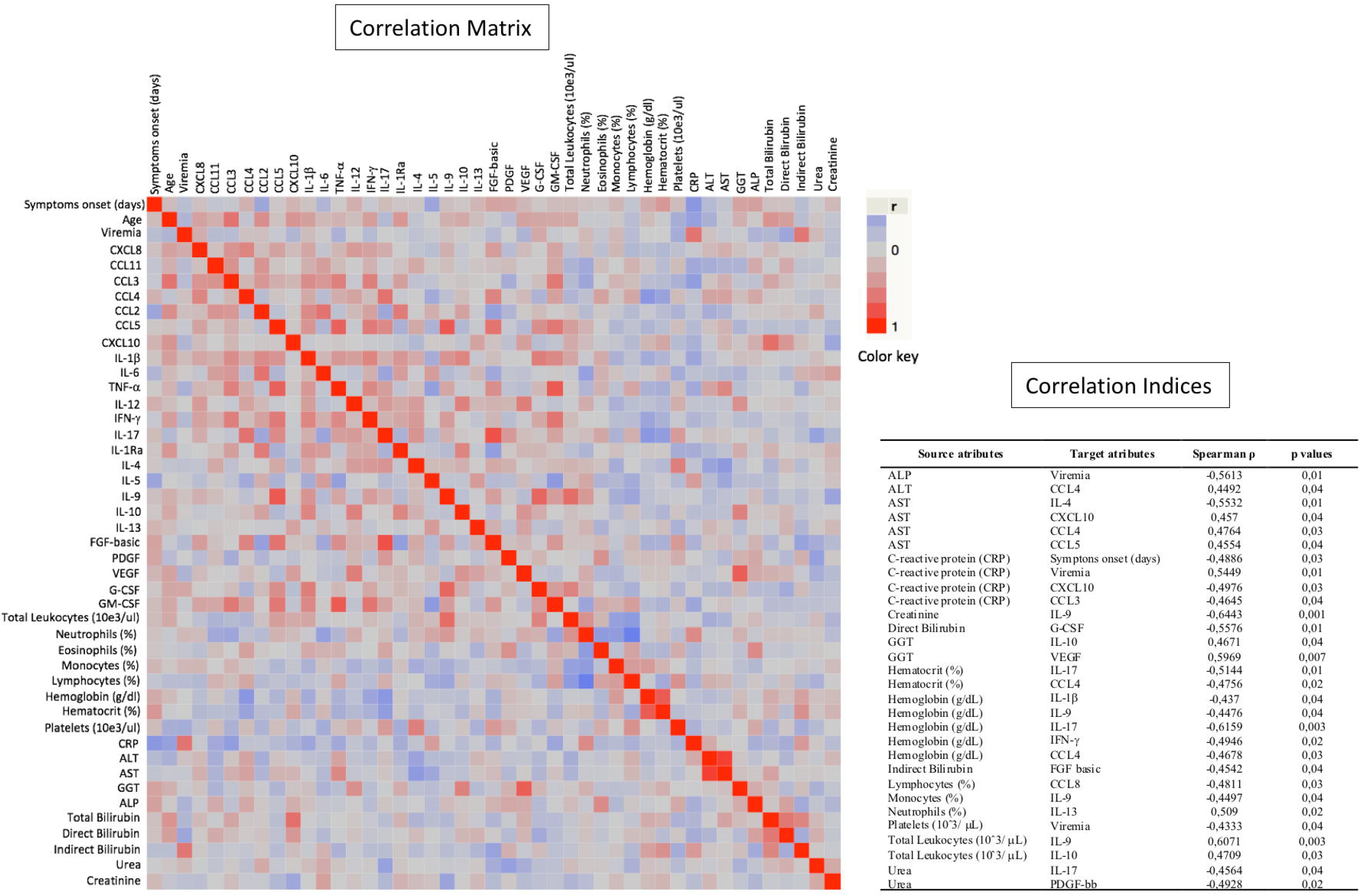
Immunological biomarkers correlations with the results of routine laboratorial tests, age, viremia, and symptoms. The nonparametric Spearman’s test was applied to evaluate multiple correlations between immunological biomarkers and the results of routine laboratorial tests. A color map matrix was plotted showing the strength and direction of these correlations (-1 blue, to +1 red). Strongly supported correlations (p < 0.05) between immunological biomarkers and routine tests are highlighted in the inserted table.

### ZIKV-infected patients display high levels of circulating biomarkers

Elevated levels of pro-inflammatory cytokines (IL-1β, IL-6, TNF-α, IFN-γ and IL-17, except IL-12 that was higher in controls), chemokines (CXCL8, CCL11, CCL3, CCL4, CCL2, CCL5 and CXCL10) and growth factors (FGF-basic, PDGF, VEGF, G-CSF and GM-CSF) were found in ZIKV-infected cases (Fig 2, light gray panels), whereas higher levels of IL-5 and IL-13 in controls (Fig 2, dark gray panels). Interestingly, the levels of IL-4 and IL-1Ra were also higher among patients. No differences were observed for the IL-9 and IL-10 (Fig 2, white panels). A similar pattern was observed when results were stratified by gender, although infected males presented significant lower levels of CCL3, CCL4, CCL5, IL-17, FGF-basic and GM-CSF. No significant differences were observed between female and male controls (Table 2).

**Fig 2.**
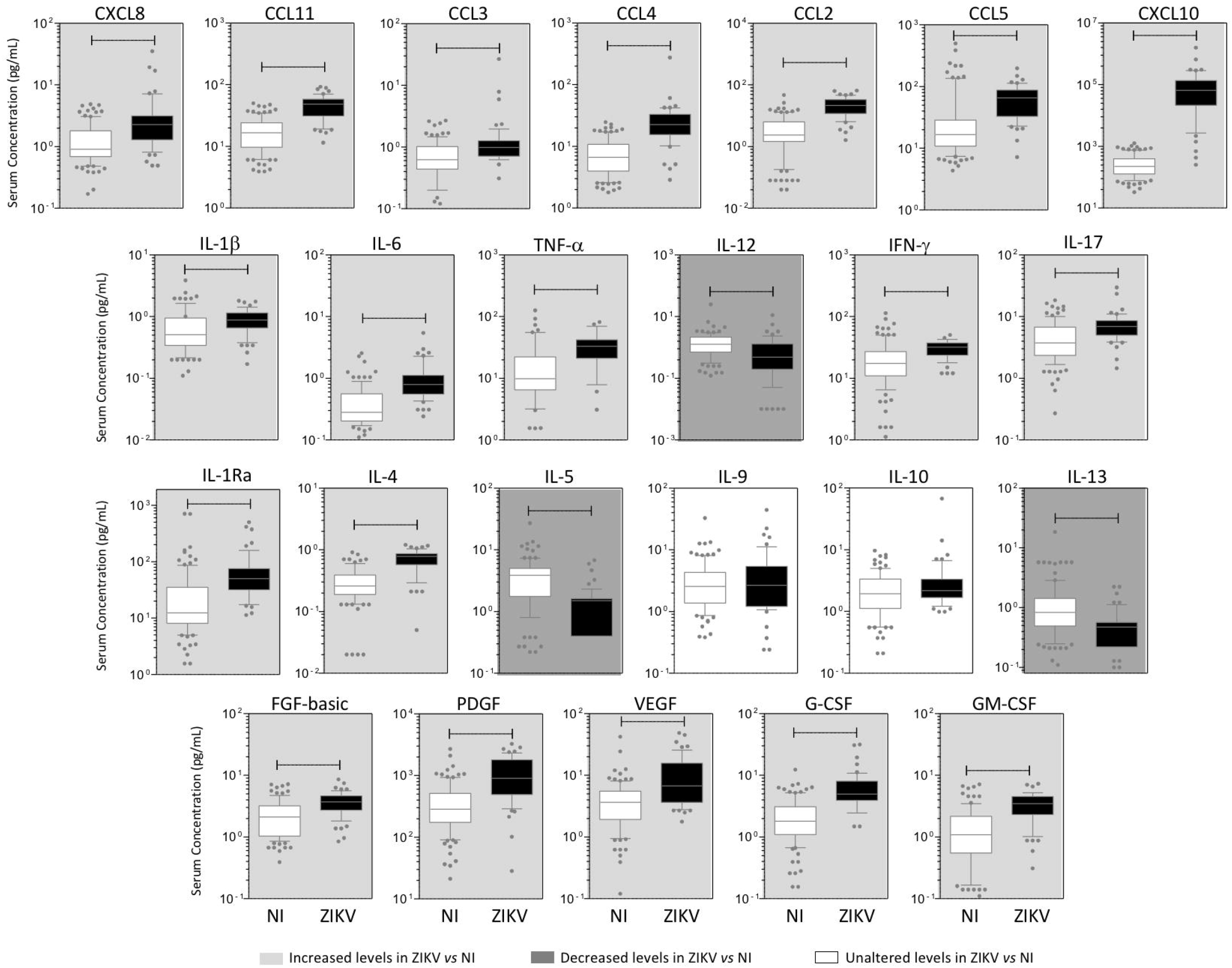
Panoramic Overview of Serum Chemokines, Cytokines and Growth Factors Early After Zika Virus Infection in Adults. Serum biomarkers (CXCL8, CCL11, CCL3, CCL4, CCL2, CCL5, CXCL10, IL-1β, IL-6, TNF-α, IL-12, IFN-γ, IL-17, IL-1Ra, IL-4, IL-5, IL-9, IL-10, IL-13, FGF-basic, PDGF, VEGF, G-CSF and GM-CSF) were measured in Zika virus-infected patients (ZIKV= 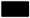, n=54) and non-infected subjects (NI= 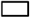, n=100) by high performance Luminex 27-plex assay as described in methods. Data expressed as pg/mL are displayed in box and whiskers (10-90 percentile) plots. Comparative analysis between NI *vs.* ZIKV was performed by Mann-Whitney test and significant differences at p<0.05 underscored by connecting lines. Colored backgrounds highlighted increased (light gray), decreased (dark gray) and unaltered (white) levels of serum biomarkers in ZIKV as compared to NI.

**Table 2.** Serum Chemokines, Cytokines and Growth Factors Early After Zika Virus Infection in Adult Females and Males.

| Analytes | Females (F), n=29 |  | *p* (1) | Males (M), n=25 |  | *p* (2) | *p* (3) | Score ZIKV/NI |  |
| --- | --- | --- | --- | --- | --- | --- | --- | --- | --- |
|  | NI | ZIKV |  | NI | ZIKV |  |  | (F) | (M) |
| **CXCL8** | 0.87<br>(0.54-1.67) | 2.26<br>(1.57-3.26) | **0.0001** | 0.98<br>(0.70-1.93) | 2.30<br>(1.19-3.11) | **0.0018** | 0.5669 | 2.6 | 2.3 |
| **CCL11** | 16.44<br>(9.29-22.22) | 48.94<br>(30.12-61.91) | **0.0001** | 16.65<br>(10.62-25.20) | 43.82<br>(29.11-56.95) | **0.0001** | 0.3488 | 3.0 | 2.6 |
| **CCL3** | 0.59<br>(0.41-0.86) | 1.15<br>(0.80-1.32) | **0.0001** | 0.65<br>(0.45-1.13) | 0.89<br>(0.67-1.07) | **0.0469** | **0.0205** | 1.9 | 1.4 |
| **CCL4** | 7.28<br>(4.56-12.20) | 28.76<br>(18.76-35.67) | **0.0001** | 5.94<br>(3.75-9.79) | 20.18<br>(12.63-26.64) | **0.0001** | **0.0162** | 4.0 | 3.4 |
| **CCL2** | 2.08<br>(1.00-4.97) | 20.73<br>(13.45-34.70) | **0.0001** | 2.46<br>(1.94-7.15) | 21.98<br>(11.86-32.34) | **0.0001** | 0.9862 | 10.0 | 8.9 |
| **CCL5** | 15.17<br>(11.36-34.98) | 82.77<br>(64.75-108.00) | **0.0001** | 17.00<br>(9.83-25.70) | 34.06<br>(23.66-65.81) | **0.0001** | **0.0001** | 5.5 | 2.0 |
| **CXCL10** | 232<br>(128-434) | 71,219<br>(32,899-148,407) | **0.0001** | 218<br>(109-392) | 44,645<br>(10,423-69,757) | **0.0001** | 0.1030 | 307 | 205 |
| **IL-1β** | 0.52<br>(0.24-0.96) | 0.93<br>(0.56-1.19) | **0.0176** | 0.52<br>(0.29-1.00) | 0.77<br>(0.61-1.14) | 0.1511 | 0.5374 | 1.8 | 1.5 |
| **IL-6** | 0.29<br>(0.20-0.57) | 0.79<br>(0.63-1.00) | **0.0001** | 0.28<br>(0.21-0.55) | 0.81<br>(0.52-1.73) | **0.0001** | 0.1944 | 2.7 | 2.9 |
| **TNF-α** | 9.76<br>(6.10-20.20) | 35.87<br>(25.45-44.08) | **0.0001** | 10.08<br>(6.45-22.62) | 26.93<br>(15.02-41.84) | **0.0015** | 0.1101 | 3.7 | 2.7 |
| **IL-12** | 1.26<br>(0.51-2.30) | 0.63<br>(0.22-1.66) | 0.0554 | 1.39<br>(0.96-2.17) | 0.34<br>(0.09-1.08) | 0.0001 | 0.1895 | 0.5 | 0.2 |
| **IFN-γ** | 14.97<br>(9.63-24.54) | 31.91<br>(26.39-38.65) | **0.0001** | 19.79<br>(14.14-27.31) | 26.41<br>(23.61-35.97) | **0.0035** | 0.4283 | 2.1 | 1.3 |
| **IL-17** | 3.84<br>(2.21-7.53) | 7.88<br>(6.28-9.02) | **0.0001** | 3.57<br>(2.50-6.76) | 5.96<br>(4.41-7.15) | **0.0114** | **0.0092** | 2.1 | 1.7 |
| **IL-1Ra** | 11.81<br>(7.81-28.66) | 47.00<br>(34.03-65.59) | **0.0001** | 12.33<br>(8.95-36.13) | 54.94<br>(30.77-115.10) | **0.0001** | 0.3623 | 4.0 | 4.5 |
| **IL-4** | 0.27<br>(0.20-0.39) | 0.81<br>(0.59-0.86) | **0.0001** | 0.26<br>(0.17-0.41) | 0.73<br>(0.53-0.90) | **0.0001** | 0.6890 | 3.0 | 2.8 |
| **IL-5** | 3.16<br>(1.67-5.02) | 1.50<br>(0.40-1.61) | **0.0001** | 4.70<br>(2.00-5.35) | 1.38<br>(1.07-1.61) | **0.0001** | 0.5792 | 0.5 | 0.3 |
| IL-9 | 2.21<br>(1.19-4.11) | 3.10<br>(1.69-5.72) | 0.0783 | 2.62<br>(1.59-4.69) | 1.42<br>(1.18-3.83) | 0.0837 | 0.0518 | 1.4 | 0.5 |
| IL-10 | 1.73<br>(0.75-3.39) | 2.11<br>(1.65-3.22) | 0.1046 | 1.95<br>(1.51-3.02) | 2.25<br>(1.52-3.41) | 0.5478 | 0.9571 | 1.2 | 1.2 |
| IL-13 | 0.75<br>(0.37-1.34) | 0.48<br>(0.22-0.57) | **0.0135** | 0.98<br>(0.80-1.57) | 0.57<br>(0.37-0.57) | **0.0001** | 0.3634 | 0.6 | 0.6 |
| FGF-basic | 1.84<br>(1.01-3.14) | <u>4.34</u><br>(3.71-5.27) | **0.0001** | 2.24<br>(1.28-3.73) | <u>3.24</u><br>(2.13-4.24) | **0.0468** | **<u>0.0014</u>** | 2.4 | 1.4 |
| PDGF | 359<br>(125-585) | 1,012<br>(616-1,933) | **0.0001** | 258<br>(196-403) | 823<br>(416-1,578) | **0.0001** | 0.3670 | 2.8 | 3.2 |
| VEGF | 2.87<br>(1.78-6.29) | 6.72<br>(4.18-16.33) | **0.0001** | 3.70<br>(2.17-4.97) | 6.26<br>(3.66-15.21) | **0.0005** | 0.5668 | 2.3 | 1.7 |
| G-CSF | 1.86<br>(1.04-2.86) | 5.96<br>(4.43-8.05) | **0.0001** | 1.83<br>(1.19-3.29) | 4.94<br>(3.42-7.66) | **0.0001** | 0.2400 | 3.2 | 2.7 |
| GM-CSF | 0.87<br>(0.52-2.00) | <u>3.76</u><br>(3.03-4.64) | **0.0001** | 1.35<br>(0.54-2.16) | <u>2.88</u><br>(1.73-3.91) | **0.0003** | **<u>0.0256</u>** | 4.4 | 2.1 |

Data are reported as median levels (IQR) in pg/mL. Statistical analysis was performed by Mann-Whitney test and significance reported as p(1), p(2) and p(3) values for comparisons between NI vs ZIKV females, NI vs ZIKV males and ZIKV females vs ZIKV males, respectively. Significant differences between NI vs ZIKV are underscored in **bold** format. Differences between ZIKV females vs ZIKV males are highlighted by **bold-underlined** format. No significant differences were observed between NI females vs NI males. Score represents the fold change (analyte median value in infected patient divided by analyte median value in controls) segregated by gender.

### Bimodal viremia is accompanied by increased levels of a defined group of biomarkers

Viremia and biomarkers were assessed at different time-points (Day 1 post-infection denoted as D1 etc.) D1 (n=11); D2 (n=13); D3 (n=10); D4 (n=09) and D5 (n=05). A bimodal distribution was observed, with two viremia peaks at D2 and D4, reaching the lowest levels at D5 (Fig 3, gray panel). Dynamics of CCL5, TNF-α, IFN-γ, IL-17 and G-CSF were closely related to viremia (Fig 3A). A similar bimodal distribution was observed for IL-1β and IL-13 (Fig 3B). The highest levels of CXCL8 and CCL2 were observed at D1 and D2 (Fig 3C). An inverse correlation was observed for IL-12, IL-10 and VEGF (Fig 3D), where the highest levels coincide with the lowest viremias. The levels of CCL3, CXCL10, IL-6 and FGF-basic display a distinct pattern, with the lowest levels observed at D3, coinciding with the first drop of viremia (Fig 3E). A valley at D4 followed by an increase at D5 was observed for CCL11, CCL4, IL-1Ra, and IL-4 (Fig 3F), and unique patterns were observed for IL-5, IL-9, PDGF, and GM-CSF (Fig 3G).

**Fig 3.**
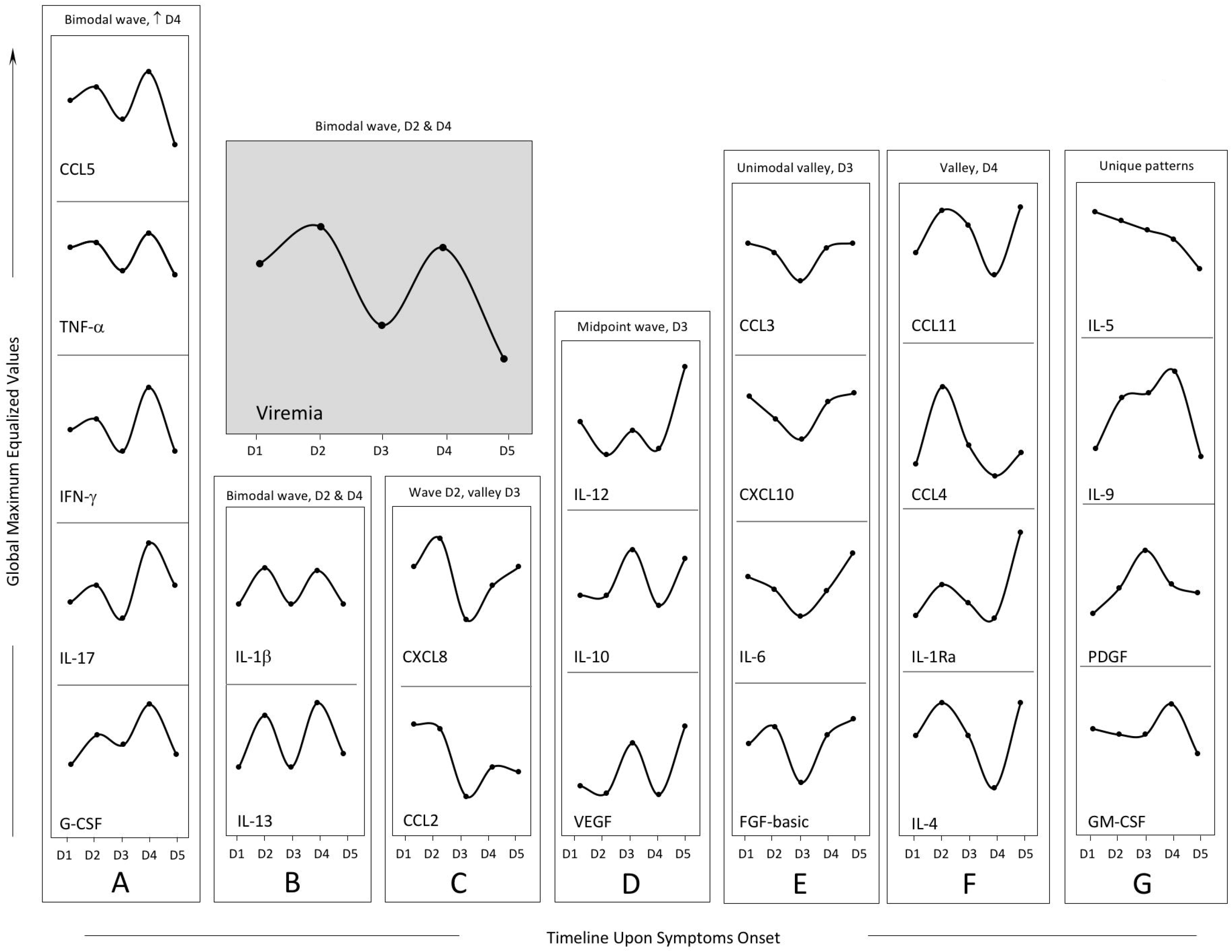
Rhythms of Viremia, Chemokines, Cytokines and Growth Factors Early After Zika Virus Infection in Adults. Cross-sectional follow-up of viremia and serum biomarkers was carried out in Zika virus-infected patients categorized according to the time (days) upon symptoms onset (D1, n=11; D2, n=13; D3, n=10; D4 n=09 and D5 n=05). Viremia (1/CT *x* 100) displayed a bimodal profile with similar waves at D2 and D4 (gray panel). Distinct patterns were identified for clusters of biomarkers as they displayed kinetic curves shaping a bimodal wave at D2 and higher wave [↑] at D4 (panel A, for CCL5, TNF-α, IFN-γ, IL-17 and G-CSF); a bimodal profile with similar waves at D2 and D4 (panel B, IL-1β and IL-13); a wave at D2 and a valley at D3 (panel C, CXCL8 and CCL2); a midpoint wave at D3 (panel D, IL-12, IL-10 and VEGF); an unimodal valley at D3 (panel E, CCL3, CXCL10, IL-6 and FGF-basic); a valley at D4 (panel F, CCL11, CCL4, IL-1Ra and IL-4) or an unique pattern (panel G, IL-5, IL-9, PDGF and GM-CSF). Data are displayed as global maximum equalized median values of the serum concentrations (pg/mL) for each biomarker.

Biomarkers were also evaluated in controls, and the IQR are represented by dashed lines (Fig 4). Most of biomarkers’ levels differ between patients and controls at all time-points, except for IL-10 at D1 and D2, IL-1β at D3. No differences were observed for IL-9.

**Fig 4.**
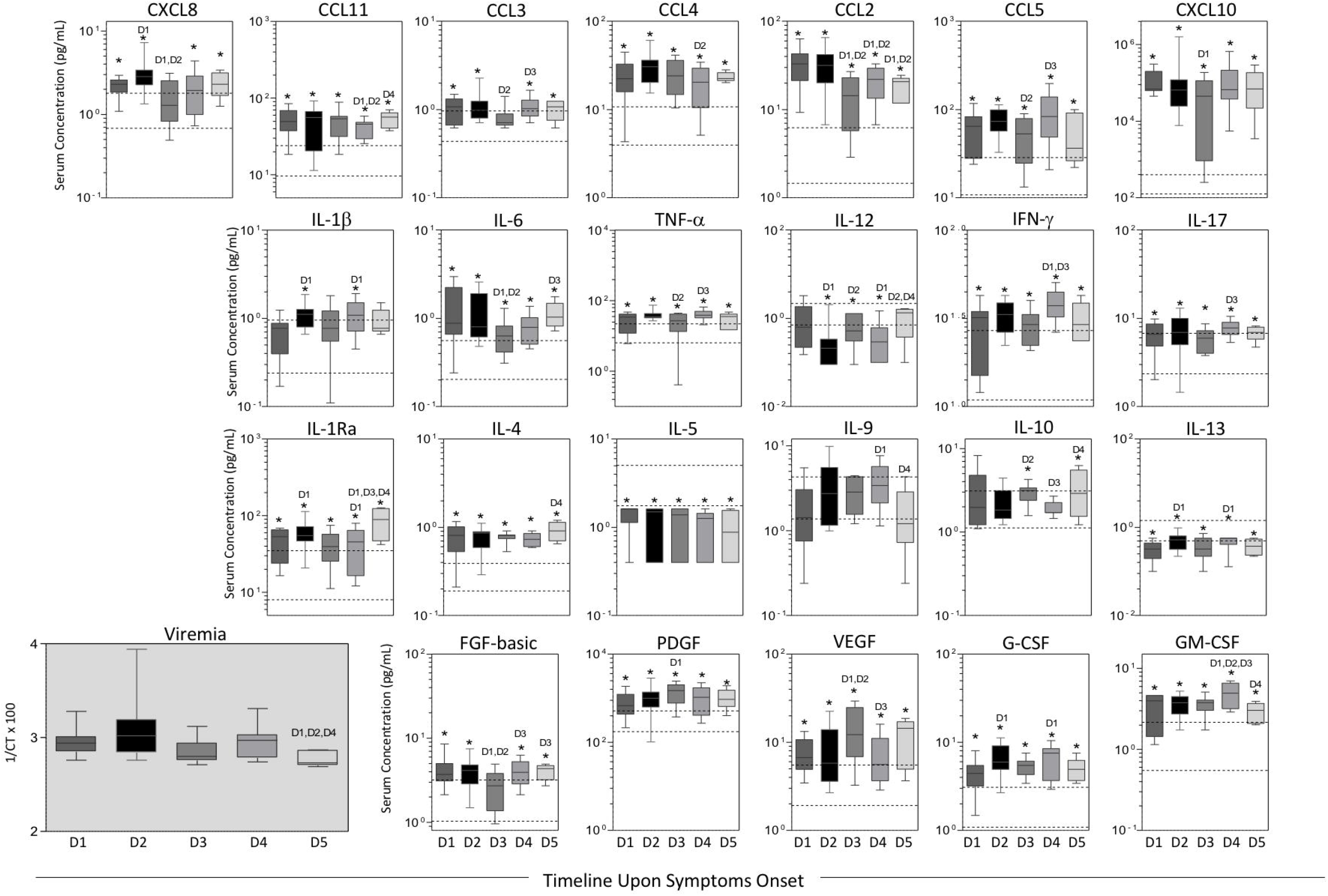
Kinetics of Viremia, Serum Chemokines, Cytokines and Growth Factors Early After Zika Virus Infection in Adults. Cross-sectional analysis of viremia and serum biomarkers was performed in Zika virus-infected patients categorized according to the time (days) upon symptoms onset (D1, n=11; D2, n=13; D3, n=10; D4 n=09 and D5 n=05). Data expressed as pg/mL are displayed in box and whiskers (10-90 percentile) plots. Multiple comparisons amongst distinct time-points upon symptoms onset were performed by Kruskal-Wallis followed by Dunn’s post-test and significant differences at p<0.05 underscored by D1, D2, D3 and D4 as they correspond to specific time-points. Comparative analysis with non-infected controls (NI) was also carried out at each time-point by Mann-Whitney test and significant differences at p<0.05 underscored by asterisks (*). Reference ranges for each biomarker were established as interquartile ranges (25^th^-75^th^ percentiles) observed in NI (dashed lines). Distinct patterns were identified for clusters of biomarkers as they displayed kinetic curves shaping a bimodal wave at D2 and higher wave [↑] at D4 (CCL-5, TNF-α, IFN-γ, IL-17 and G-CSF); a bimodal profile with similar waves at D2 and D4 (IL-1β and IL-13); a wave at D2 and a valley at D3 (CXCL8 and CCL2); a midpoint wave at D3 (IL-12, IL-10 and VEGF); an unimodal valley at D3 (CCL3, CXCL10, IL-6 and FGF-basic); a valley at D4 (CCL11, CCL4, IL-1Ra and IL-4) or an unique pattern (IL-5, IL-9, PDGF and GM-CSF).

### ZIKV infection elicited a set of general and timeline-specific biomarkers

The biomarker levels were used to build a signature (Fig 5, left panels) as described in the methods section. A significant difference in the overall profile was observed in ZIKV-infected cases (Fig 5, top-left panel). Furthermore, the radar chart revealed that 19/24 (79%) biomarkers were highly induced by ZIKV infection (Fig 5, bottom-left panel). Almost all biomarkers analyzed were found in levels above the global median in more than 75% of the infected patients.

**Fig 5.**
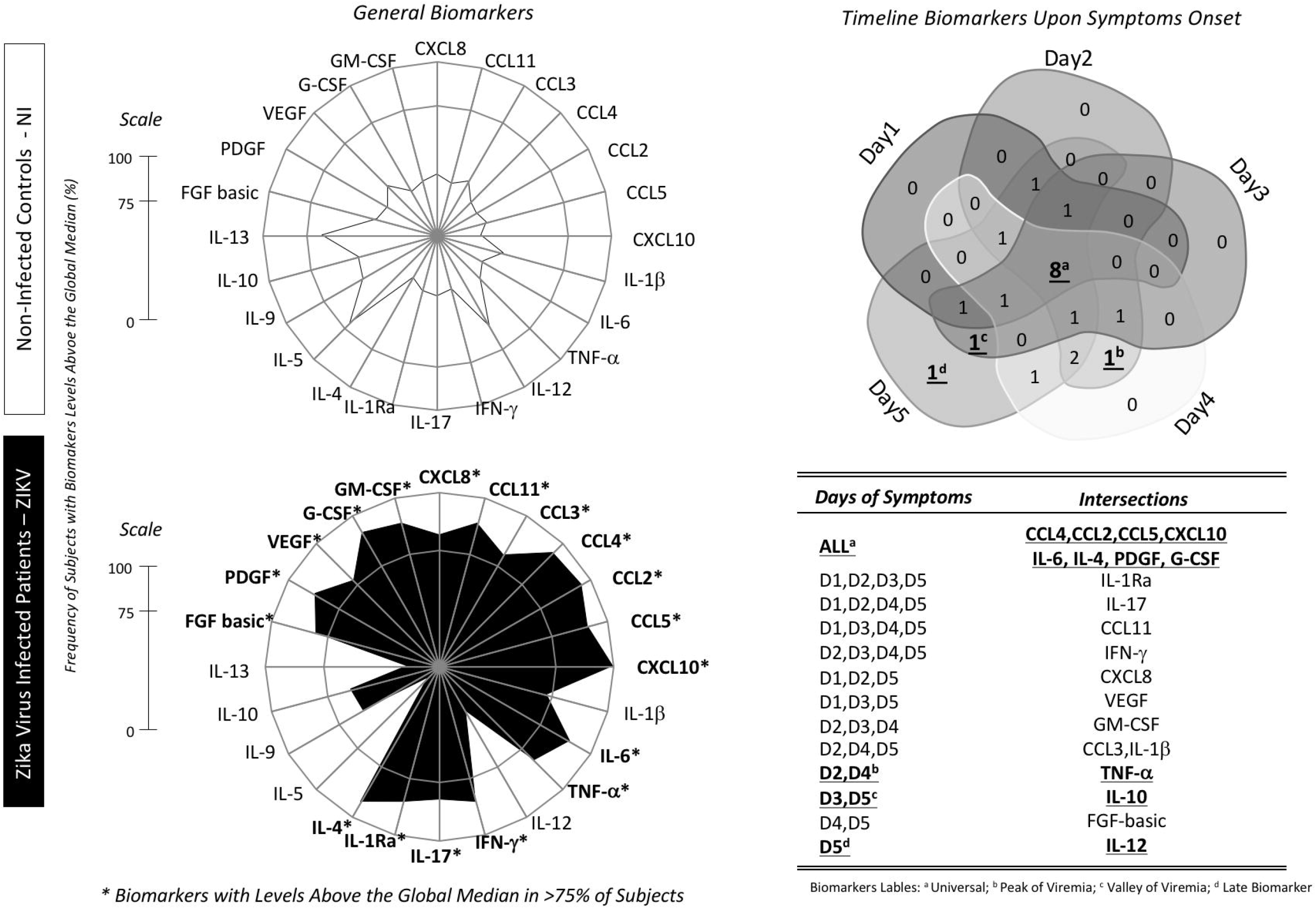
General and Timeline Biomarkers upon Symptoms Onset Early After Zika Virus Infection in Adults. Biomarker signatures of NI (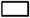) and ZIKV (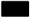) were constructed as described in methods. Data are presented in radar charts as the proportion of subjects with serum biomarker levels above the global population median values (NI plus ZIKV). Biomarkers with levels above the global median in more than 75% of subjects were highlighted by asterisks (*). The Venn diagram shows the intersections with common attributes as well as selective biomarkers along the timeline of symptoms onset (Day 1; Day 2; Day 3; Day 4 and Day 5). Venn diagram report summarizes selected attributes with patterns labeled as (a) universal; (b) peak of viremia; (c) valley of viremia or (d) late biomarkers (inserted table).

Venn diagram analysis showed that four chemokines (CCL4, CCL2, CCL5, CXCL10); two cytokines (IL-6, IL-4) and two growth factors (PDGF, G-CSF) were significantly induced in all time-points (Fig 5, right panel). Of note, TNF-α appears as a single biomarker at the intersection of the viremia peaks (D2 and D4). In contrast, IL-10 is the only unregulated biomarker at viremia valleys (D3 and D5) while increased levels of IL-12 appears at D5 (Fig 5, inserted table).

### Distinct biomarker networks are observed at different time-points

Cytoscape software was used to assemble correlative analysis of immunological biomarkers. The exploratory analysis demonstrated that earlier infection was associated with more imbricate and complex biomarker networks. Most correlations at D1 and all correlations at D2 were positive (solid lines). The level of complexity decreased from D1 to D5. However, the interactions were more complex at D2 and D4, concomitant with the viremia peaks (Fig 6).

**Fig 6.**
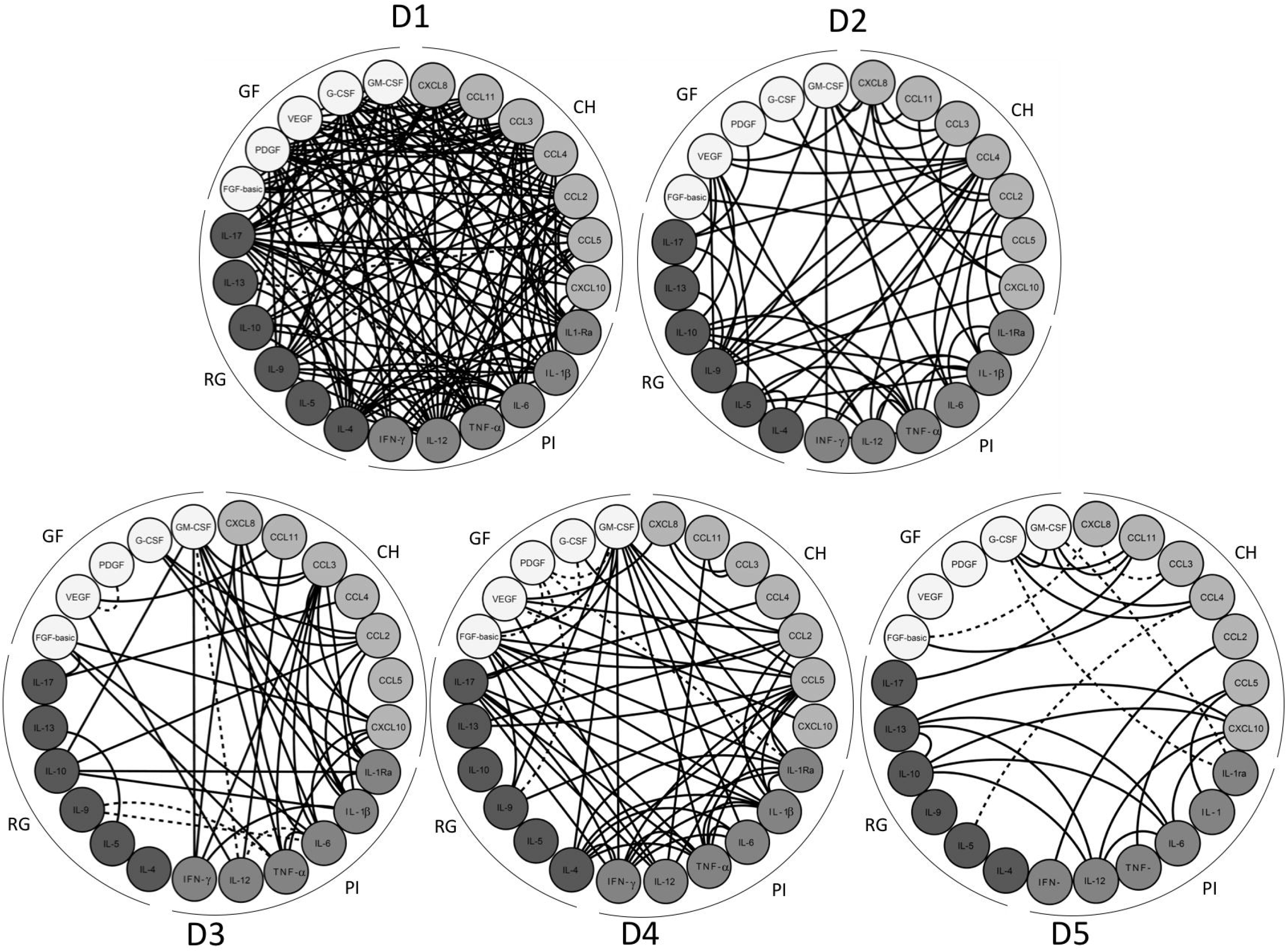
Timeline Biomarker Networks Early After Zika Virus Infection in Adults. Systems integrative biology analysis of attributes was assembled using Cytoscape software platform to build circular nodal network layout for each time-point upon Zika virus infection day 1 (D1) up day 5 (D5) based on Spearman’s correlation matrices. Significance was considered at p<0.05. The timeline of networks is displayed as circular layouts to characterize the interaction along the early time-points. Colored nodes were employed to identify chemokines (CH), pro-inflammatory cytokines (PI), regulatory cytokines (RG) and growth factors (GF). Connecting edges were drawn to underscore the association between attributes, classified as positive (solid line) or negative (dashed line).

### High-dimensional data analysis elected CXCL10 as the most promising biomarker for a putative clinical application

A heatmap matrix was constructed to evaluate the profile of biomarkers associated with ZIKV infection. This analysis demonstrated that CXCL10 clustered most patients, segregating them from controls. Additionally, a decision tree was built to identify the biomarker most able to segregate patients. This approach confirmed the heatmap observations indicating CXCL10 as the most relevant element, followed by IL-4 and VEGF. The analysis showed a very high global accuracy (99.4%) with a leave-one-out-cross-validation of 96.8% (Fig 7). The significance of these attributes (CXCL10, IL-4 and VEGF) was assessed by 3D-plots and the performance of the root attribute (CXCL10) evaluated by scatter plot distribution and ROC curve analysis (Fig 7, bottom panels). CXCL10 alone lead to a very high global accuracy ranging from 0.952-0.998. Together, the results demonstrated that CXCL10 measurement ascertains 94% of the patients, with no false-positive identification and outstanding indices (co-positivity, conegativity and likelihood ratio).

**Fig 7.**
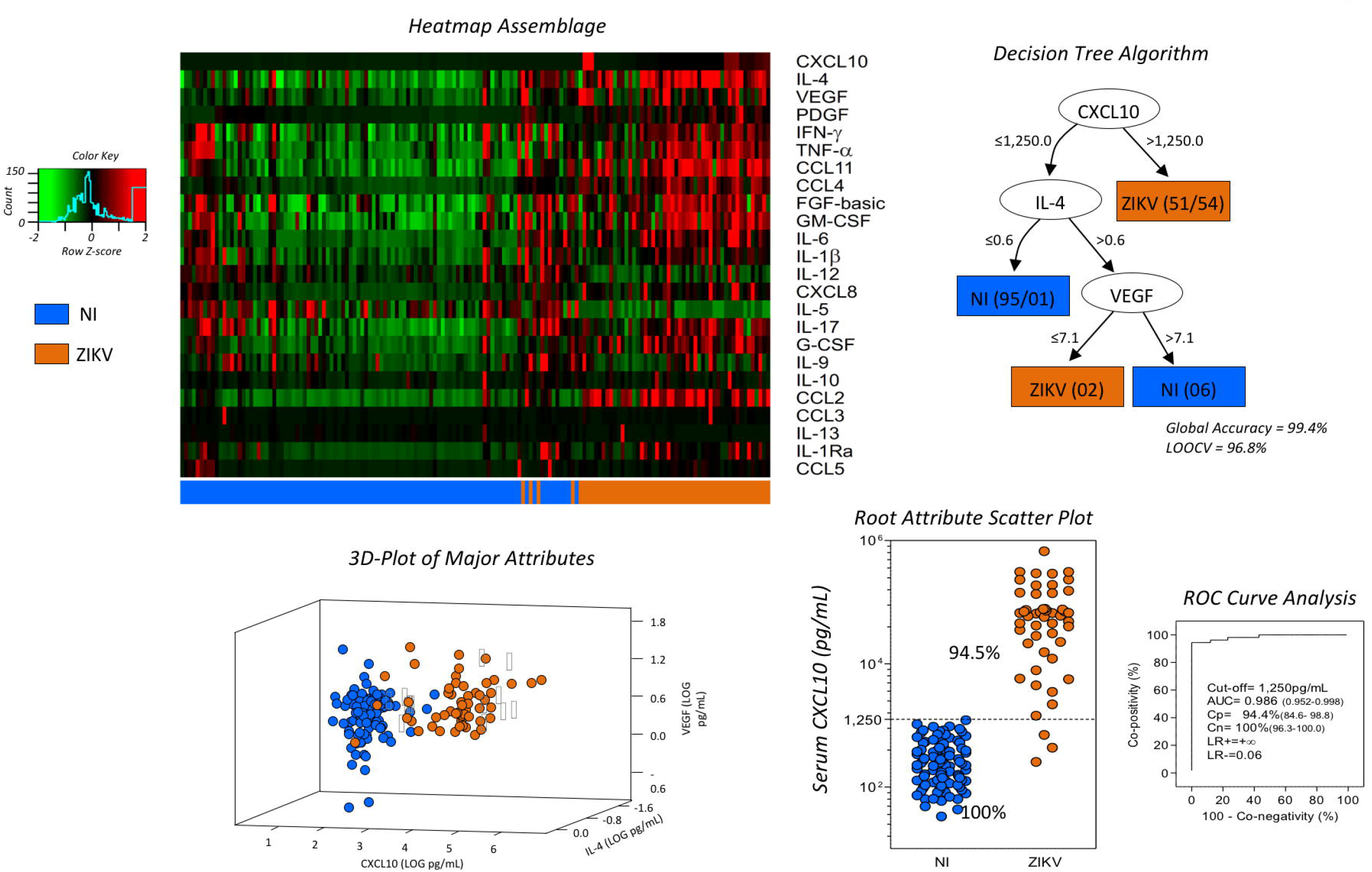
High-dimensional Data Analysis Early After Zika Virus Infection in Adults. Machine-learning high-dimensional data approaches were applied to further explore and identify feasible criteria applicable for the clinical follow-up of Zika virus infection. (A) Heatmap panels were built to verify the ability of attributes to segregate ZIKV (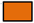) and NI (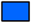) groups as they present low (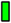) or high (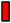) levels of serum biomarkers. Decision tree algorithms were generated define root and branch attributes to segregate patients (ZIKV=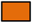) from non-infected controls (NI=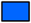). Global accuracy and leave-one-out-cross-validation (LOOCV) values are provided in the figure. The root/branch attributes selected by the decision tree algorithm were compiled into a 3D-plot to verify their clusterization strength. The performance of the selected root attribute to discriminate ZIKV (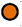) from NI (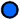) was evaluated by scatter plot distribution and validated by receiver operating-characteristic indices (Area under the curve, AUC; Co-positivity, Cp; Co-negativity, Cn; Positive/Negative Likelihood Ratio, LR+/LR-).

## Discussion

The pathogenesis of ZIKV infection is still largely unknown, and the main determinants of disease manifestations are not yet well established. Understanding serum immunomodulators during acute infection may be a first step to elucidate the mechanisms underlying ZIKV-induced immunopathology.

We show that the immune response during the acute phase of ZIKV infection is polyfunctional and broadly inflammatory as evidenced by significant elevated levels of IL-4, IL-17, IFN-γ, IL-1β, IL-1Ra, TNF-α and IL-6 in patients. This is consistent with findings from Kam et al [9] that also found a robust pro-inflammatory cytokine response during acute ZIKV infection with elevations of IL-18, TNF-α, IFN-γ, IL-8, IL-6, GRO-α, and IL-7. Alternatively, when we stratified the results by gender, ZIKV-infected males presented lower levels of CCL3, CCL4, CCL5, IL-17, FGF-basic and GM-CSF. The reason for this difference is unknown in the context of ZIKV infection, however this finding is consistent with the literature demonstrating that females tend to mount a higher innate and adaptive immune system response to viruses compared to men [14]. In addition, this result may also be explained as females were sampled on average one day earlier than men with the median time from onset to diagnostic sampling (Females= day 2; Males= day 3).

It is possible that previous exposure to Flavivirus antigens may affect the immune response to ZIKV infection. In the present study, almost all patients (51/54) exhibited positive DENV IgG antibodies. Manaus has had several dengue epidemics, including co-circulation of different serotypes [15-17]. Moreover, the Amazonas State is endemic for Yellow Fever virus (YFV) and has a very high YFV-vaccination coverage. Thus, mostly individuals enrolled in this study have experienced previous Flavivirus exposure potentially modulating the cytokine and chemokine responses. These differences in prior Flavivirus exposure may account for some differences in cytokine and chemokine profiles shown by Kam et al. that examined a Brazilian cohort of patients from Campinas, Brazil where Yellow Fever vaccination was not required by the government at the time of their study as it is in Manaus, Brazil.

Similar to our results, comparable immune response induced during the acute phase has previously been described in infections caused by ZIKV and other Flaviviruses, including YFV, DENV and West Nile virus [8,18-20]. In the case of ZIKV infection, the mechanism of inflammatory immune response is not clearly delineated. The immune response may be triggered by viral upregulation of expression of pattern recognition receptors (PRRs) engaged in downstream pathways and inflammatory antiviral response such as IRF7, IFN-α, IFN-β, and CCL5 [7]. Interestingly, we showed a strong positive correlation between IFN-α and CCL5, suggesting that the synergistic effect of these cytokines might be crucial on the outcomes of the acute inflammation caused by ZIKV.

Our findings also revealed higher levels of growth factors and chemokines among patients. Likewise, increased levels of CXCL10, CCL5, CCL3 and VEGF were primarily demonstrated in patients acutely infected with ZIKV, while elevated levels of GM-CSF, CCL4, and FGF-basic biomarkers only in the recovery phase [8]. Our study demonstrated that all chemokines and growth factors analyzed were significantly increased in the acute phase in comparison with non-infected controls. In fact, the role of growth factors in the pathogenesis of arboviruses infections remains a matter of debate [20-22]. We demonstrate that a remarkable increase of FGF-basic, PDGF, VEGF, G-CSF and GM-CSF identifies the acute phase of ZIKV infection, which suggests the importance of chemokines and growth factors in the initiation and regulation of the acute-phase immune response.

Similarly, increased serum concentrations of both CXCL (CXCL8 and CXCL10) and CCL chemokines (CCL2, CCL3, CCL4, CCL5, and CCL11) were found in acute ZIKV infection. The role of CCL5 in the arbovirus-induced immunopathology remains a controversial issue, but this chemokine along with CCL2 and CCL3 were previously linked to severity of dengue and Japanese encephalitis virus infections, including neurological diseases and impairment of neuronal survival [23-27].

Furthermore, we found strong correlations between TNF-α and CCL5 concentrations and percentages of circulating neutrophils and lymphocytes in acute ZIKV infection. This finding is likely due to the role of TNF-α and CCL5 in leukocyte chemo attraction [28,29] and demonstrates the important role of this cytokine and chemokine in stimulation of the innate and adaptive immune system in response to ZIKV infection. In addition, this manuscript is the first to describe the bimodal nature of viremia in acute Zika infection and corresponding peaks in inflammatory cytokine production. A biological model explaining bimodal viremia was firstly described in the classical study of Fenner’s with the Mousepox virus [30]. Similarly, Flaviviruses are initially replicated in Langerhans cells at the site of inoculation and in draining regional lymph nodes. Despite a robust anti-viral innate immune response that eliminates viral infected cells, some virus particles are disseminated by blood (primary viremia). Therefore, several organs and tissues may become infected producing a second wave of viral replication that reaches blood causing a secondary viremia [31]. The equine infection by African Horse Sickness Viruses, another arbovirus of the *Orbivirus* genus, *Reoviridae* family, also shows two viremia peaks. The first peak is observed after the viral multiplication into lymph nodes, whereas the second peak is observed after viral replication in spleen, lungs and endothelial cells [32]. Several arboviruses are known to cause prolonged viremia into their natural hosts, and this is well documented for encephalitic Alphaviruses [33,34] leading to higher transmission rates for mosquito vectors. Interestingly, bimodal viremia has been found in patients after low dose live attenuated 17DD Yellow Fever vaccine administration [35]. The low dose live attenuated vaccine is hypothesized to elicit a less robust immune response in comparison to the standard dosage vaccine that does not clear the initial viremia leading to a second peak of viremia a few days later. Further research to determine if ZIKV undergoes similar processes is needed.

In this manuscript, we reported high levels of pro-inflammatory mediators during the acute phase of ZIKV infection. Paradoxically, although the inflammatory response leads to viral clearance, the high levels of circulating pro-inflammatory biomarkers may facilitate the transmission of viruses from circulation to the central nervous system by increasing the permeability of the blood brain barrier. This phenomenon has been already reported for the West Nile virus [36], another neurovirulent Flavivirus, and may partially explain ZIKV neuroinvasiveness.

Remarkably, the CXCL10 was expressed greater than 200-fold in ZIKV-infected subjects. Augmented serum levels of CXCL10 have been found during severe clinical manifestations of dengue and Yellow fever [14,18,37]. CXCL10 has also been shown to play an important role in CD-8+ T-cell recruitment as part of an anti-flaviviral response in the central nervous system to West Nile virus [38] and dengue virus [39]. Furthermore, CXCL10 has been previously associated as a biomarker of severity in several diseases including those caused by bacteria like *Mycobacterium tuberculosis* and *Legionella pneumophila*; protozoans like *Trypanosoma brucei, Leishmania major, Plasmodium vivax* or *Plasmodium falciparum* [40]; viral diseases such as in Simian Human Immunodeficiency Virus Encephalitis [41] and viral acute respiratory infection in healthy adults, mainly those caused by Influenza virus [42].

Of paramount importance, CXCL10 overexpression has been observed in non-infectious neuronal diseases like Alzheimer’s and multiple sclerosis, and in infectious diseases like HIV-associated dementia [43]. Furthermore, different studies showed that the over-expression of CXCL10 leads to apoptosis in fetal neurons [40] that is triggered by intracellular Ca(2+) elevation activating caspase-9 and caspase-3 [43]. CXCL10 has also been strongly implicated in Guillain-Barré syndrome pathogenesis [44]. Thus, we hypothesize that the high elevations of CXCL10 in ZIKV patients may contribute to neuronal damage affecting the developing fetal brain and potentially targeting peripheral nerves in Guillain-Barré syndrome as well. Consistent with this hypothesis, Kam et al. specifically identified higher levels of CXCL10 in ZIKV-infected patients with neurological complications compared to those without and higher levels of CXCL10 in ZIKV-infected pregnant women carrying babies with fetal growth associated malformations.

High levels of CXCL10 have been previously described at acute and convalescent phases, with more prominent expression at the latter [8]. Unfortunately, although our data strongly suggest CXCL10 as a biomarker of ZIKV acute infection, we were unable to perform a longitudinal analysis to verify its kinetics across different stages of the disease, to further confirm whether the concentrations of this chemokine would be down or up-regulated. In addition, CXCL10 elevation is also observed in pre-eclampsia and hypertension found in pregnancy resulting in a range of fetal injuries, including intrauterine growth retardation and neurological damage induced by hypoxia [45,46]. Thus, it is reasonable to suggest that ZIKV-induced inflammation may increase fetal injuries.

CXCL10 may also be an important therapeutic target [40]. For example, CXCL10 neutralization by specific antibodies or genetic deletion in CXCL10-/-mice protected against cerebral malaria infection and inflammation [47]. Passive transfer of anti-CXCL10 antibodies reduced inflammatory leukocyte recruitment across the blood brain barrier. Furthermore, statin medications commonly used for cholesterol control have been shown to decrease CXCL10 and to be effective in CXCL10 mediated Crohn’s disease [48].

Finally, we describe the relationship between the timing of viremia and cytokine elevations. The acute phase of ZIKV infection lasts around five days [49]. This study assessed the acute phase biomarkers and viral titers at different time-points (until day 5). Augmented levels of CCL4, CCL2, CCL5, CXCL5, CXCL10, IL-6, IL-4, PDGF, and G-CSF immunomodulators were observed at all time-points. The peak of viremia, at Day 2 and Day 4, was accompanied by increased TNF-α levels. Instead, the IL-10 elevation appeared to be directly related to the lowest virus titers (Day 3 and Day 5), while the highest levels of IL-12 were found at Day 5. These findings allow us to deduce that the acute phase of ZIKV is characterized mainly by an innate immune system inflammatory response, with the overlap of the inflammatory biomarkers and viremia peaks, and anti-inflammatory response coinciding with viremia decay.

Altogether, this study identifies unique characteristics of the acute inflammatory and multifactorial immune response induced by ZIKV and depicts CXCL10 as a potential biomarker of the acute infection, perhaps, a predictor of severity. Nevertheless, further longitudinal studies that measure the host immunopathological aspects at several time-points is required to better characterize all the immunological factors involved in the Zika disease. The elevated concentrations of serum biomarkers observed in this study, may bring new insights to the ZIKV immunopathology puzzle.

## Acknowledgments

The authors are thankful to the Programa de Desenvolvimento Tecnológico em Insumos para a Saúde - PDTIS-FIOCRUZ for the use the flow cytometry (CPqRR) and RealTime PCR (ILMD) facilities.

